# PTEN and the PTEN-like phosphatase CnrN have both distinct and overlapping roles in a *Dictyostelium* chemorepulsion pathway

**DOI:** 10.1101/2024.02.23.581751

**Authors:** Kristen M. Consalvo, Ramesh Rijal, Steven L. Beruvides, Ryan Mitchell, Karissa Beauchemin, Danni Collins, Jack Scoggin, Jerome Scott, Richard H. Gomer

## Abstract

The directed movement of eukaryotic cells is crucial for processes such as embryogenesis and immune cell trafficking. The enzyme Phosphatase and tensin homolog (PTEN) dephosphorylates phosphatidylinositol 3,4,5-trisphosphate [PI(3,4,5)P_3_] to phosphatidylinositol 4,5-bisphosphate [PI(4,5)P_2_]. *Dictyostelium discoideum* cells require both PTEN and the PTEN-like phosphatase CnrN to locally inhibit Ras activation to induce biased movement of cells away from the secreted chemorepellent protein AprA. Both PTEN and CnrN decrease basal levels of PI(3,4,5)P_3_ and increase basal numbers of macropinosomes, and AprA prevents this increase. AprA requires both PTEN and CnrN to increase PI(4,5)P_2_ levels, decrease PI(3,4,5)P_3_ levels, inhibit proliferation, decrease myosin II phosphorylation, and increase filopod sizes. AprA causes PTEN, similar to CnrN, to localize to the side of the cell towards AprA in an AprA gradient. However, PTEN and CnrN also have distinct roles in some signaling pathways. PTEN, but not CnrN, decreases basal levels of PI(4,5)P_2_, AprA requires PTEN, but not CnrN, to induce cell roundness, and CnrN and PTEN have different effects on the number of filopods and pseudopods, and the sizes of filopods. Together, our results suggest that CnrN and PTEN play unique roles in *D. discoideum* signaling pathways, and possibly dephosphorylate PI(3,4,5)P_3_ in different membrane domains, to mediate chemorepulsion away from AprA.

## Introduction

The directed movement of eukaryotic cells toward or away from an external stimulus is crucial for neuronal migration, embryogenesis, and the trafficking of immune cells during inflammation (Borrell, 2019; Garcia-Cuesta et al., 2019; Glass et al., 2020; SenGupta et al., 2021). Studies on the movement of the model eukaryote *Dictyostelium discoideum* have revealed mechanisms of chemoattraction, where cells move toward a stimulus. Migrating *D. discoideum* cells extend pseudopods to adhere to the substrate and contract the trailing edge of the cells to force them forward (Uchida et al., 2003; Van Haastert and Devreotes, 2004).

During development, a gradient of the chemoattractant cyclic adenosine monophosphate (cAMP) activates phosphatidylinositolL3LOH kinases (PI(3)Ks) that increases levels of phosphatidylinositol 3,4,5-trisphosphate [PI(3,4,5)P_3_] and active Ras at the *D. discoideum* cell membrane to form polymerized actin-rich pseudopods projecting towards the source of cAMP (Cheng et al., 2020; Chung et al., 2000; Huang et al., 2003; Kortholt et al., 2013; Park et al., 2004; Sasaki et al., 2004; Zigmond et al., 1997). *D. discoideum* cells utilize Phosphatase and tensin homolog (PTEN) to dephosphorylate phosphatidylinositol 3,4,5-trisphosphate [PI(3,4,5)P_3_] to phosphatidylinositol 4,5-bisphosphate [PI(4,5)P_2_], which prevents PI(3,4,5)P_3_-dependent filamentous actin (F-actin) accumulation and pseudopod formation at the side of the cell facing away from the source of cAMP, causing biased movement of cells toward cAMP (Chung et al., 2000; Funamoto et al., 2002; Huang et al., 2003; Iijima and Devreotes, 2002; Park et al., 2004; Sasaki et al., 2004; Wessels et al., 2007; Zigmond et al., 1997). Also at the side of the cell facing away from the source of cAMP, polymerized myosin II provides contractile force for the cell movement (Egelhoff et al., 1993; Heissler and Sellers, 2016; Levi et al., 2002; Liang et al., 1999; Pasternak et al., 1989). The PTEN-like phosphatase cell number regulator N (CnrN) is a *D. discoideum* PTEN-like protein with phosphatidylinositol phosphatase activity, sharing 23-25% sequence identity with PTENs, including *D. discoideum* PTEN (Tang and Gomer, 2008). Loss of CnrN in *D. discoideum* cells leads to an increase in cAMP production during development (Tang and Gomer, 2008). The loss of PTEN or CnrN causes increased PI(3,4,5)P_3_ accumulation, and the loss of PTEN increases F-actin levels after cAMP stimulation (Huang et al., 2003; Iijima and Devreotes, 2002; Tang and Gomer, 2008).

Proliferating *D. discoideum* cells secrete a protein called autocrine proliferation repressor protein A (AprA) (Brock and Gomer, 2005; Choe et al., 2009; Phillips and Gomer, 2012). AprA inhibits *D. discoideum* proliferation (Brock and Gomer, 2005). In addition, AprA is a chemorepellent for *D. discoideum* cells, causing them to move in a biased direction away from a source of AprA (Phillips and Gomer, 2012). At In a colony of cells, there will be a high extracellular concentration of AprA at the center, and low concentrations or AprA outside the colony, causing a gradient of AprA at the edge of the colony (Gomer, 2019; Kirolos and Gomer, 2022; Rijal et al., 2019). This gradient of a chemorepellent causes cells at the colony edge to move away from the colony, potentially in search of new food sources (Phillips and Gomer, 2012). In a gradient of AprA, the side of the cell sensing the highest concentration of AprA shows an inhibition of Ras activation, F-actin formation, and pseudopod formation, inhibiting the movement of cells toward the higher concentration of AprA, and thus biased movement of cells away from the higher concentration of AprA (Kirolos and Gomer, 2022; Rijal et al., 2019).

In a uniformly high concentration of extracellular AprA, the AprA inhibition of Ras activation and pseudopod formation occurs on all sides of a cell, and this causes cells to round up and stop moving (Kirolos and Gomer, 2022). At the center of a large colony of densely populated cells, the cells have overgrown the local food source and no longer need to move around and hunt for food. In this region of a colony, the AprA concentration is high and relatively uniform (Kirolos and Gomer, 2022). Presumably to conserve energy, the high AprA concentrations causes cells to stop moving and become round (Kirolos and Gomer, 2022; Phillips and Gomer, 2012).

Although PTEN and CnrN both dephosphorylate PI(3,4,5)P_3_ to PI(4,5)P_2_, and would thus appear to be redundant, AprA-induced chemorepulsion requires both of these enzymes (Herlihy et al., 2013; Phillips and Gomer, 2014; Rijal et al., 2019; Tang et al., 2018). In this report, we elucidate the overlapping and different roles of PTEN and CnrN in the AprA signal transduction pathway.

## Results

### PTEN is localized to the side of the cell towards AprA in an AprA gradient

*Dictyostelium* cells exposed to a gradient of the chemorepellent AprA exhibit a biased movement away from AprA (Phillips and Gomer, 2012). Both PTEN and the PTEN-like phosphatase CnrN are required for AprA-induced chemorepulsion (Herlihy et al., 2013; Rijal et al., 2019). Expressing PTEN-GFP in *pten^–^* cells (*pten^–^/pten-gfp*) (Iijima and Devreotes, 2002) (confirmed by PCR using gene-specific primers, **Supplementary Figures 1A and B**) restored the ability of cells to move away from AprA (**Supplementary Figure 2A**). This suggests that the insensitivity to AprA observed in *pten^–^* cells is specifically due to the loss of PTEN, and not influenced by secondary mutations. Consistent with previous observations in wild-type Ax2, *cnrN^–^*, and *pten^–^* cells (Rijal et al., 2019), AprA did not affect the speed and the persistence of *pten^–^/pten-gfp* cells (**Supplementary Figure 2B and C**). CnrN localizes to the side of the cell facing AprA in an AprA gradient (Rijal et al., 2019). We observed a uniform distribution of PTEN at the cell periphery in unstimulated cells, and in an AprA gradient, the percentage of cells with PTEN at the side facing AprA increased, while the percentage of cells with PTEN at the side facing away from AprA decreased (**Supplementary Figure 2D and E**). Together, these data indicate that in a gradient of AprA, like CnrN, PTEN tends to localize to the side of the cell towards the source of AprA.

*PTEN and CnrN have different effects on the accumulation of PI(4,5)P_2_ and PI(3,4,5)P*_3_ *during chemorepulsion*

PTEN dephosphorylates PI(3,4,5)P_3_ to PI(4,5)P_2_ at the side of *Dictyostelium* cells facing away from cAMP, inhibiting the formation of filamentous actin (F-actin) and preventing pseudopod formation (Funamoto et al., 2002; Iijima and Devreotes, 2002; Matsuoka and Ueda, 2018; Sasaki et al., 2007; Wessels et al., 2007). Loss of either PTEN or CnrN in *Dictyostelium* cells causes overaccumulation of PIP3 in response to cAMP stimulation (Iijima and Devreotes, 2002; Tang and Gomer, 2008). To determine how two enzymes with redundant functions, PTEN and CnrN, regulate PI(4,5)P_2_ and PI(3,4,5)P_3_ levels during chemorepulsion, Ax2, *pten^–^*, *cnrN^–^*, *pten^–^/pten-gfp*, or *cnrN^–^/cnrN-gfp* cells (verified by PCR using gene-specific primers, **Supplementary Figure 1A and B**) were exposed to a uniform concentration of AprA and phosphoinositides were measured. The basal levels of PI(4,5)P_2_ were higher in *pten^–^* and *pten^–^/pten-gfp* cells than in Ax2, *cnrN^–^,* or *cnrN^–^/cnrN-gfp* cells (**Figure 1A**). Both *pten^–^* and *cnrN^–^* cells had increased basal PI(3,4,5)P_3_ levels compared to Ax2 (**Figure 1B**). Expressing *pten-gfp* in *pten^–^* cells or expressing *cnrN-gfp* in *cnrN^–^* cells reduced PI(3,4,5)P_3_ levels (**Figure 1B**). AprA increased PI(4,5)P_2_ levels at 5 and 10 minutes in Ax2 cells (**Figure 1C**). In *pten^–^* cells, AprA reduced PI(4,5)P_2_ levels at 5, 10, and 20 minutes (**Figure 1D**). Expression of PTEN-GFP in *pten^–^* cells, as with Ax2 cells, caused AprA to increase PI(4,5)P_2_ levels at 10 minutes (**Figure 1E**). AprA did not significantly affect PI(4,5)P_2_ levels at any time in *cnrN^–^* cells, but as with Ax2 cells, increased PI(4,5)P_2_ levels in *cnrN^–^/cnrN-gfp* cells at 10 minutes (**Figure 1F and G**).

**Figure 1.**
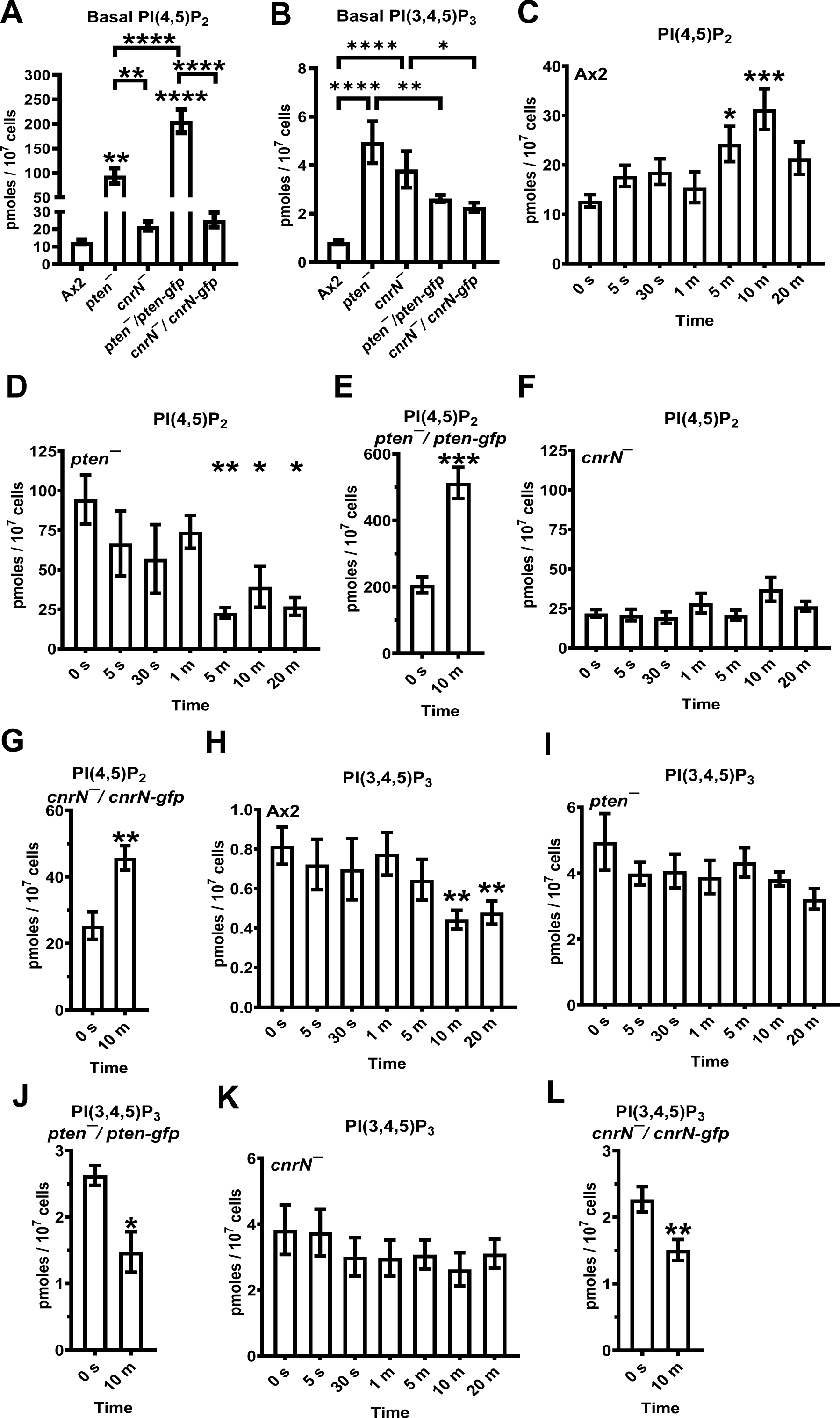
PTEN and CnrN cause differential effects on PI(4,5)P_2_ and PI(3,4,5)P_3_ accumulation during chemorepulsion. (A-L) Cells were incubated in HL5 with 300 ng/ml AprA or an equivalent volume of buffer for the indicated amounts of time. Phosphatidylinositol were extracted and quantified using PI(4,5)P_2_ (A, and C-G) and PI(3,4,5)P_3_ (B, and H-L) ELISAs. Values in (A) and (B) are the 0 seconds values from (C - L). Values are mean ± SEM from 4 (C, J, and L), 5 (D, E, G, I, and K), or 6 (F, H) independent experiments. * p L 0.05, ** p L 0.01, *** p L 0.001, **** p L 0.0001 compared to the WT (A and B) or the 0 time point (C-L), unless otherwise indicated (A and B, One-way ANOVA with Tukey’s correction) (C-L, Mann-Whitney *U* test).

AprA caused a decrease in PI(3,4,5)P_3_ levels at 10 and 20 minutes in Ax2 cells (**Figure 1H**). In *pten^–^* cells, AprA did not affect PI(3,4,5)P_3_ levels, but as with Ax2 cells, AprA reduced PI(3,4,5)P_3_ levels in *pten^–^/pten-gfp* cells at 10 minutes (**Figure 1I and J**). Similarly, AprA did not affect PI(3,4,5)P_3_ levels in *cnrN^–^* cells but reduced PI(3,4,5)P_3_ levels in *pten^–^/pten-gfp* cells at 10 minutes (**Figure 1K and L**). Together, these data indicate that PTEN but not CnrN decreases basal levels of PI(4,5)P_2_, that (assuming that the PTEN levels in *pten^–^/pten-gfp* are not exactly the same as in Ax2 cells) the proper levels of PTEN are needed to maintain basal levels of PI(4,5)P_2_, that both PTEN and CnrN decrease basal levels of PI(3,4,5)P_3_, that AprA increases levels of PI(4,5)P_2_ and that CnrN is needed for this, that in the absence of PTEN but not CnrN AprA causes an unknown enzyme to decrease levels of PI(4,5)P_2_, that AprA decreases PI(3,4,5)P_3_ levels, and that both PTEN and CnrN are needed for this effect.

### AprA requires CnrN and PTEN to inhibit proliferation

AprA inhibits the proliferation of wild-type *Dictyostelium* cells, and the loss of CnrN abolishes this effect (Brock and Gomer, 2005; Herlihy et al., 2013). To determine whether AprA requires PTEN to inhibit proliferation, proliferating Ax2, *cnrN*^–^, and *pten*^–^ cells were incubated with AprA, and the decrease in cell density compared to a buffer control after a 24-hour incubation was determined. As previously observed, AprA inhibited the proliferation of Ax2 cells (Brock and Gomer, 2005) (**Figure S2F**). Both *cnrN^–^* and *pten*^–^ cells had a slower proliferation than Ax2, and AprA did not significantly affect their proliferation (**Figure S2F**). Together, these data suggest that AprA requires both CnrN and PTEN to inhibit proliferation.

### AprA requires CnrN and PTEN to decrease myosin II phosphorylation

In a cAMP gradient, a localized increase in PI(3,4,5)P_3_ levels at the side of the cell closest to the source of cAMP increases actin polymerization and pseudopod formation, causing biased movement of cells towards cAMP (Cheng et al., 2020; Chung et al., 2000; Huang et al., 2003; Park et al., 2004; Sasaki et al., 2004; Zigmond et al., 1997). Myosin II stabilizes the cytoskeleton by associating with the actin meshwork and provides force on actin filaments (Egelhoff et al., 1993; Heissler and Sellers, 2016; Levi et al., 2002; Liang et al., 1999; Pasternak et al., 1989). Myosin II is active in its filamentous form, which is negatively regulated by phosphorylation (Liang et al., 1999). Both *cnrN^–^* and *pten*^–^ cells had reduced basal total actin levels but *cnrN^–^* had normal basal total myosin II levels (**Figure 2A and B**). In contrast, *cnrN^–^* and *pten*^–^ cells had increased basal F-actin and polymerized myosin II levels (**Figure 2C and D**). Loss of CnrN but not PTEN decreased basal phosphorylated myosin levels compared to Ax2 (**Figure 2E**). Similar to its effects on Ax2 cells, AprA did not alter levels of total actin, total myosin II, F-actin, and polymerized myosin II in *pten^–^* and *cnrN^–^* cells (**Supplementary Figure 3**). AprA decreased levels of phosphorylated myosin II after 5 minutes in Ax2 cells (Rijal et al., 2019) (**Figure 2F**). Loss of CnrN or PTEN prevented that decrease in phosphorylated myosin II levels (**Figure 2G and H**). Together, these data suggest that both PTEN and CnrN are required for AprA-induced myosin II dephosphorylation.

**Figure 2.**
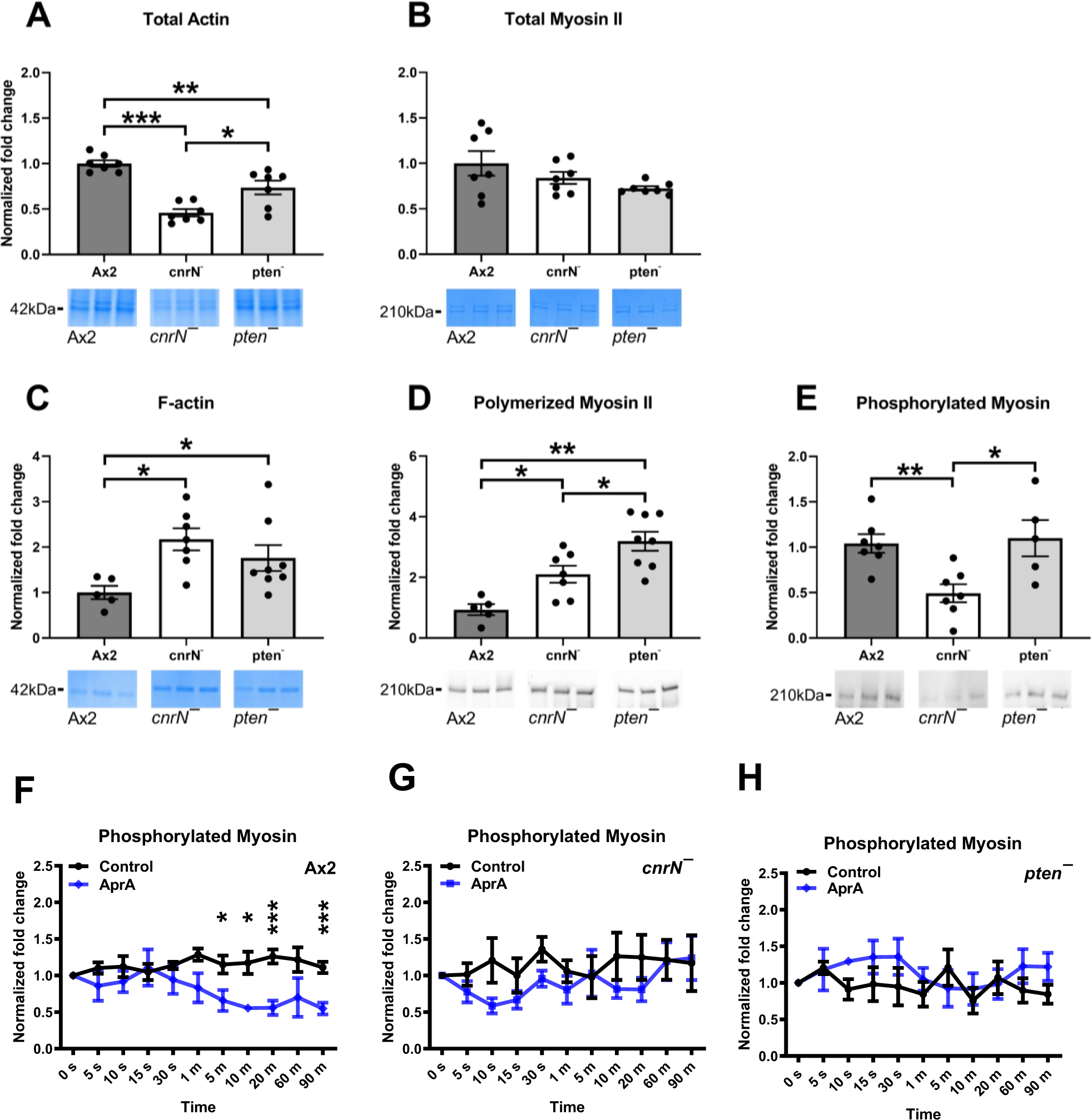
Loss of CnrN or PTEN in *Dictyostelium* cells reduces total actin and increases F-actin and polymerized myosin II levels. (A – E) The whole cell lysates (A and B) or the detergent-insoluble cytoskeletons (C - E) from Ax2, *cnrN^–^* and *pten^–^* cells were resolved by SDS-PAGE and gels were stained with Coomassie (A - C), or Western blots of gels were stained with anti-myosin II antibodies (D) or anti-P-threonine antibodies (E). Actin (A and C), myosin II (B and C), or phosphorylated myosin (E) levels were estimated by densitometric analysis. The average of total actin, polymerized myosin II, F-actin, or phosphorylated myosin from Ax2 cells was considered 1. (F – H) Cells were incubated in growth medium (0 seconds) or in growth medium with 300 ng/ml AprA for the indicated times, and phosphorylated myosin levels were estimated as described in (E). Phosphorylated myosin levels at time 0 was considered 1. Values are mean ± SEM for ≥ 3 independent experiments. * p L 0.05, *** p L 0.001 (Mann-Whitney *U* test).

### AprA requires CnrN and PTEN to inhibit Ras activation

In a cAMP gradient, the activation of Ras at the front of the cell (the side closest to the source of the cAMP attractant) leads to the localized PI3 kinase activation and PI(3,4,5)P_3_ production at the front, resulting in actin polymerization and pseudopod formation (Cheng et al., 2020). During chemorepulsion, AprA prevents pseudopod formation by inhibiting Ras activation at the side of the cells facing towards AprA, causing biased movement of cells (Kirolos and Gomer, 2022). To determine if AprA requires CnrN and PTEN to inhibit Ras activation, Ax2, *cnrN^–^*, and *pten^–^* cells were treated with AprA for 0, 10, and 30 minutes, and Ras activation was assessed using a pull-down assay of active Ras with Raf-RBD affinity beads. The loss of CnrN or PTEN did not significantly affect the basal levels of total and active Ras (**Figure 3A-C**). AprA did not significantly affect total Ras levels in *cnrN^–^* and *pten^–^* cells at 10 and 30 minutes (**Figure 3D**). As previously observed (Kirolos and Gomer, 2022), AprA reduced active Ras levels in Ax2 within 30 minutes (**Figure 3E**). The loss of CnrN or PTEN abolished this effect (**Figure 3E**). Together, these data suggest that AprA requires both CnrN and PTEN to inhibit Ras activation.

**Figure 3.**
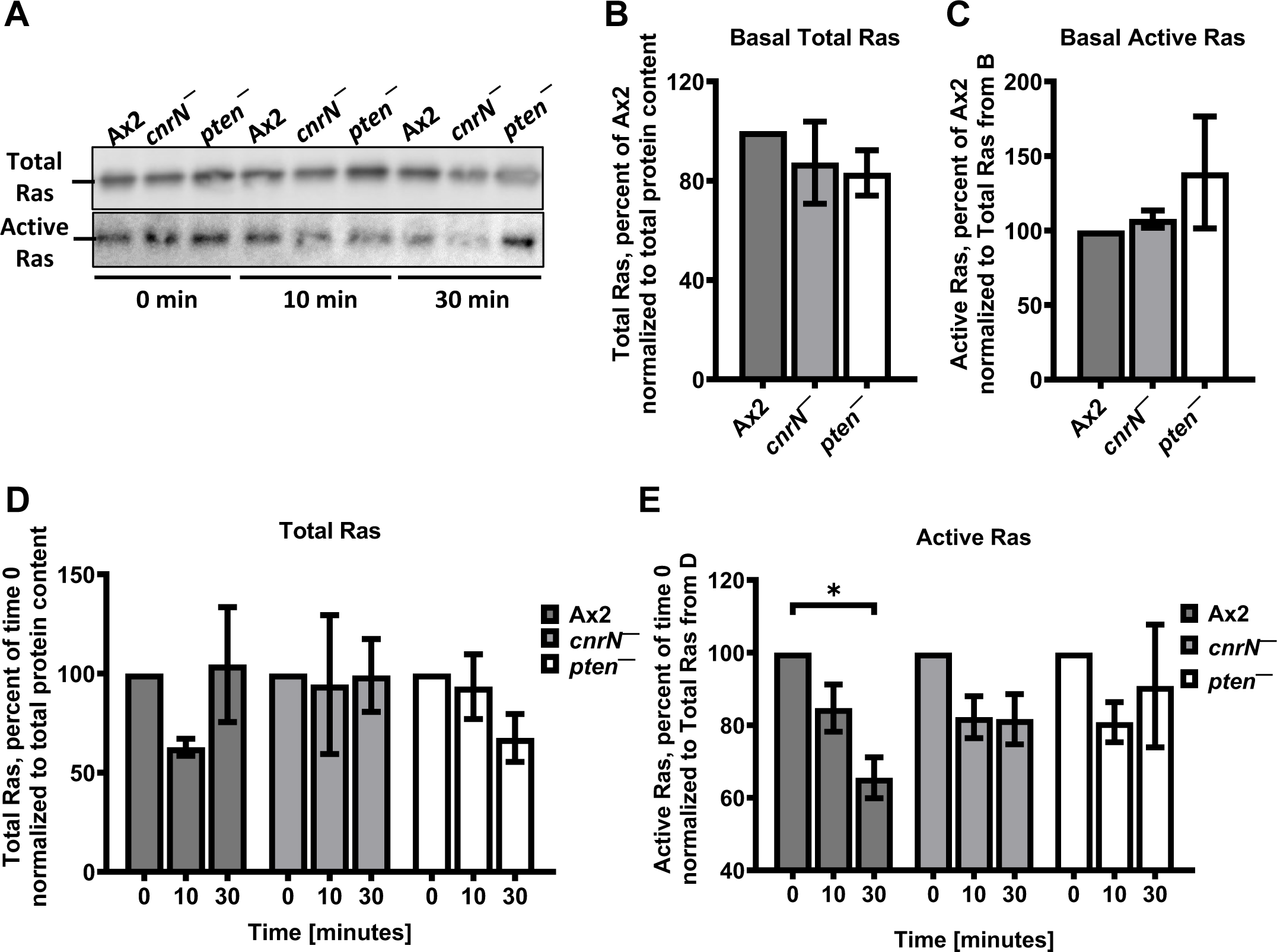
AprA inhibits Ras activation in Ax2, but not in *cnrN^–^* or *pten^–^* cells. (A) Cells of indicated strains were incubated in growth medium with 300 ng/ml AprA for indicated times, and total cell lysates or Raf-RBD affinity bead pull-down samples were run on SDS-polyacrylamide gels. Western blots of the gels were stained with anti-Pan Ras antibodies (A). Images are representative of 3 independent experiments. Densitometry was used to estimate levels of Ras in the Raf-RBD affinity bead pull-down assays (B - E). * p < 0.05 (Unpaired t test with Welch’s correction).

### AprA requires PTEN, but not CnrN, to increase the roundness of cells

In a gradient of AprA, *Dictyostelium* cells inhibit Ras activation and pseudopod formation at the side of the cell facing the source of AprA (Kirolos and Gomer, 2022; Rijal et al., 2019). Prolonged exposure (60 minutes) of cells to uniform concentration of AprA causes cells to become rounder (Kirolos and Gomer, 2022). To determine if AprA-induced cell roundness is dependent on cell density, the roundness of Ax2 cells at 1, 1.5, 5, and 10 x 10^5^ cells/ml densities was measured by determining the ratio of the short and long axes of the cell (short/long) before adding AprA, and 30 minutes after adding AprA or buffer control. Cell densities did not significantly affect the roundness of cells before adding AprA, but after further incubation for 30 minutes in the absence of AprA, cells became rounder as cell densities increased (**Figures 4A and B**). AprA further increased the roundness of cells as cell densities increased (**Figures 4A and B**). To determine if AprA requires CnrN and/or PTEN to induce cell rounding, Ax2, *cnrN^–^*, and *pten^–^* cells were treated with uniform concentration of AprA or buffer control for 30 minutes, and cell roundness was determined. AprA increased the roundness of Ax2 and *cnrN^–^* but not *pten^–^* cells (**Figure 4C and D**). Together, these data suggest that AprA induces cell roundness in a cell density-dependent and PTEN-dependent manner.

**Figure 4.**
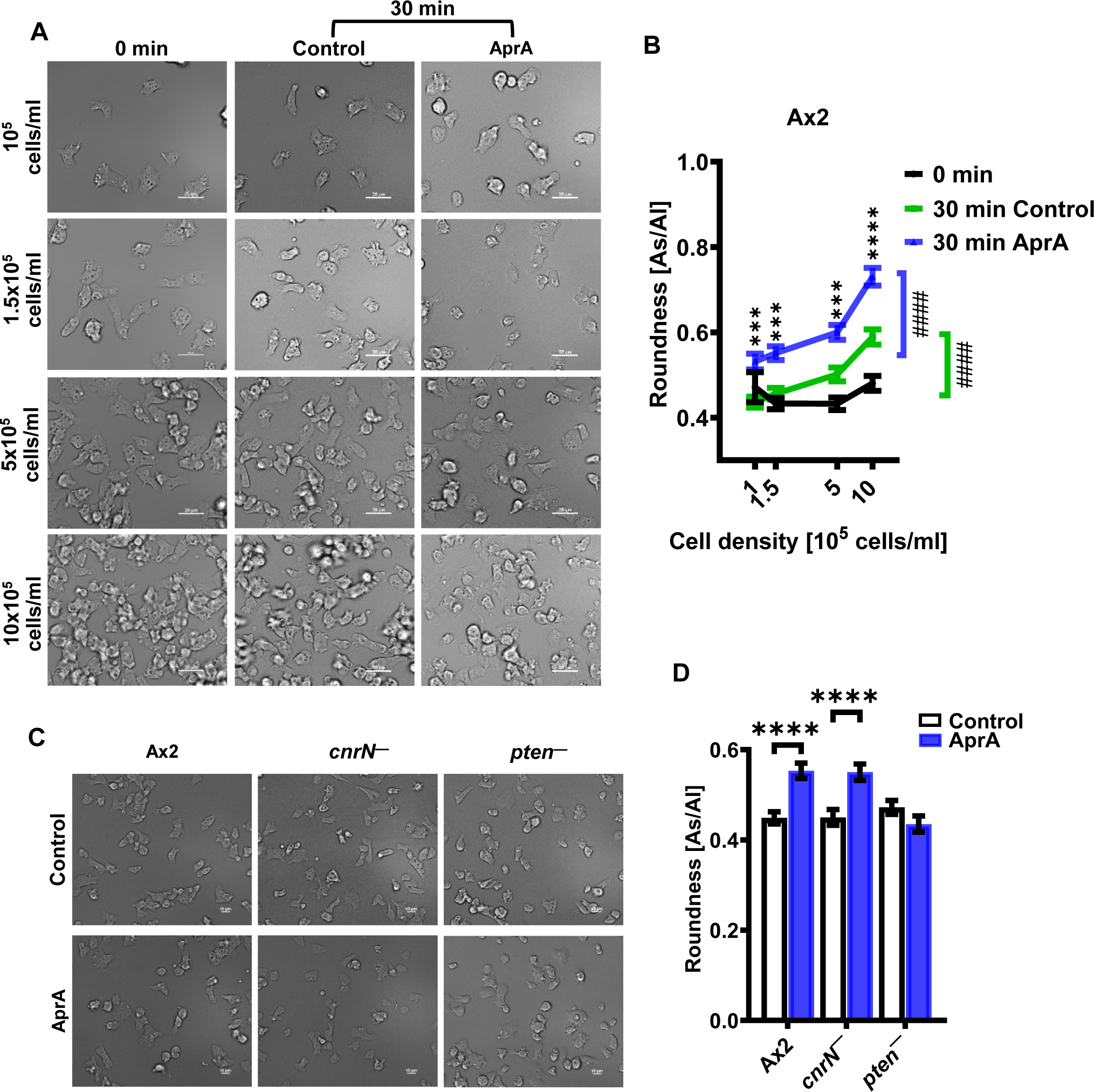
AprA causes Ax2 and *cnrN^–^* cells, but not *pten^–^* cells, to become rounder. (A and B) Ax2 cells at the indicated densities were incubated in growth medium for 30 minutes. Images of cells were captured to determine the roundness of cells at 0 minutes. Subsequently, AprA was added to a final concentration of 300 ng/ml, or an equivalent volume of buffer (control) was added to the cells. The cells were then incubated for an additional 30 minutes and images of cells were taken (A), and the roundness of cells was determined (B). Roundness was assessed by calculating the ratio of the short (As) and long (Al) axes of cells (As/Al). As/AI = 1 indicates perfectly round cells. (C and D) Cells of the indicated strains at a density of 1.5×10^5^ cells/ml were treated with 300 ng/ml AprA or an equivalent volume of buffer (control) for 30 minutes, images cells were taken (C), and roundness (D) was measured as in (A and B). Bars are 20 µm (A) or 10 µm (C). Images are representative for each experiment. Values represent the mean ± SEM of 40 cells per experiment from three independent experiments. ^####^ p < 0.0001 (compared between 1 x10^5^ and 10×10^5^ cells/ml in each condition in B; color coded for each condition), and *** p < 0.001 and **** p < 0.0001 (Two-way ANOVA with Šídák’s multiple comparisons test between control and AprA for each cell density in B and unpaired t test with Welch’s correction in D).

### AprA requires CnrN and PTEN to prevent an increase in the number of macropinosomes and filopod sizes

*Dictyostelium* cells increase formation of pseudopods and filopods at the front of the cell in a steep gradient of cAMP but do not alter pseudopod formation in a shallow gradient of cAMP (Bosgraaf and Van Haastert, 2009; Heid et al., 2005). *Dictyostelium* cells use cup-shaped ruffles to uptake liquid nutrient in a process called macropinocytosis (Williams and Kay, 2018). Fast-moving cells have slow rate of macropinocytosis, and vice versa (Veltman, 2015). An AprA gradient does not alter the rate of pseudopod formation but increases the rate of filopod projections (Rijal et al., 2019). To determine if AprA requires CnrN and PTEN to regulate membrane protrusions, Ax2, *cnrN^–^*, and *pten^–^* cells were incubated in HL5 on a coverslip-bottom petri plate in the presence or absence of a uniform concentration of AprA, and observed with an inverted microscope with an oil-immersion objective under the culture, and the microscope condenser placed in the HL5 closely over the cells. Starting at 0, 30, and 60 minutes, images of cells were taken for 5 minutes. The numbers, sizes, and lifespans of filopods, pseudopods, and macropinosomes (**Supplementary Figure 4A**) were then determined. Compared to Ax2, *cnrN^–^* cells had a reduced number of filopods, and *pten^–^* had increased filopod size and pseudopod numbers (**Supplementary Figure 4**). Compared to our previous study (Rijal et al., 2019), the size of pseudopods in Ax2 cells were approximately 10 times larger. In the previous study, we used cells taken from shaking suspension culture, and in this study, we used cells that had been growing on surfaces in stationary submerged culture. The difference may thus be due to the different culture conditions. Together, these data suggest that CnrN but not PTEN increases the number of filopods, and that PTEN but not CnrN decreases filopod sizes and the number of pseudopods.

Ax2 cells had an increased number of filopods after 60 minutes, and AprA did not significantly affect that number (**Supplementary Figure 5A**). This increase may have been due to hypoxia in the confined environment under the condenser. Compared to Ax2, AprA increased the number of filopods in *cnrN^–^* but not *pten^–^* cells over the first 5 minutes (time 0), and that effect of AprA was lost at 30 and 60 minutes (**Supplementary Figure 5B and C**). As previously observed (Rijal et al., 2019), AprA did not significantly affect numbers of pseudopods. Compared to Ax2, AprA decreased the number of pseudopods in *cnrN^–^* at 30 minutes and *pten^–^* cells at 0 and 30 minutes (**Supplementary Figure 5D – F**). In untreated *pten^–^* cells at 60 minutes, the basal number of pseudopods was reduced compared to cells at 0 and 30 minutes (**Supplementary Figure 5F**). As above, this may have been due to hypoxia. Ax2 cells increased number of macropinosomes at 30 and 60 minutes, and AprA prevented that increase (**Supplementary Figure 5G**). AprA did not significantly affect the number of macropinosomes in *cnrN^–^* and *pten^–^* cells (**Supplementary Figure 5G - I**).

AprA increased filopod size at time 0 in Ax2 cells but not in *cnrN^–^* and *pten^–^* cells (**Supplementary Figure 6A - C**). AprA did not significantly affect the sizes of pseudopods or macropinosomes in Ax2, *cnrN^–^* and *pten^–^* cells (**Supplementary Figure 6D - I**). AprA increased the lifespan of filopods and pseudopods in *pten^–^* but not Ax2 or *cnrN^–^* cells at 60 minutes and did not affect lifespan of macropinosomes in Ax2, *cnrN^–^*, and *pten^–^* cells (**Supplementary Figure 7A - I)**. Together these data suggest that for unknown reasons, AprA transiently increases the number of filopods in *cnrN^–^* cells, that AprA transiently increases the number of pseudopods and that this effect is reversed in cells lacking either CnrN or PTEN, that AprA increases the number of macropinosomes and this effect is reversed in cells lacking either PTEN or CnrN, that AprA transiently increases the filopods size and this effect requires either CnrN or PTEN, and that AprA increases lifespan of filopods and pseudopods in cells lacking PTEN.

## Discussion

PTEN and the PTEN-like phosphatase CnrN that dephosphorylate PI(3,4,5)P_3_ to PI(4,5)P_2_ are necessary for AprA-induced chemorepulsion (Herlihy et al., 2013; Phillips and Gomer, 2014; Rijal et al., 2019; Tang et al., 2018). In this report we determined how *D. discoideum* cells utilize PTEN and CnrN in response to AprA, as prevailing evolutional ideology has been that superfluous proteins and functions are lost, as seen by the disappearance of CnrN in higher level eukaryotes (Tang and Gomer, 2008). We found PTEN decreases basal levels of PI(4,5)P_2_ and PI(3,4,5)P_3_, and CnrN decreases basal levels of PI(3,4,5)P_3_. Both PTEN and CnrN are required to increase the number of macropinosomes by an unknown mechanism, and AprA prevents that increase. AprA requires both PTEN and CnrN to increase PI(4,5)P_2_ levels, decrease PI(3,4,5)P_3_ levels, inhibit proliferation, decrease myosin II phosphorylation, inhibit Ras activation, and increase in filopod sizes, but only requires PTEN to induce cell roundness.

PTEN and CnrN are cytosolic and/or uniformly distributed on the cytosolic side of the plasma membrane of unstimulated *D. discoideum* cells (Funamoto et al., 2002; Tang and Gomer, 2008). In a cAMP gradient, PTEN localizes to the side of the cell facing away from the source of cAMP, prevents PI(3,4,5)P_3_ accumulation via conversion of PI(3,4,5)P_3_ to PI(4,5)P_2_, and suppresses localized F-actin accumulation and lateral pseudopod formation (Cheng et al., 2020; Funamoto et al., 2002; Iijima and Devreotes, 2002; Matsuoka and Ueda, 2018). In an AprA gradient, similar to CnrN (Rijal et al., 2019), PTEN localize to the side of the cell facing toward the source of AprA, suggesting that these proteins functions concurrently to suppress PI(3,4,5)P_3_ dependent F-actin polymerization and pseudopod formation in the region of the cell closest to the source of AprA.

Loss of PTEN but not CnrN increased basal PI(4,5)P_2_ levels, possibly due to dysregulation of CnrN activity as a result of either loss of PTEN or overexpression of PTEN in cells lacking PTEN and optimum PTEN levels appears to be necessary to maintain basal PI(4,5)P_2_ levels. Loss of either PTEN or CnrN increased basal PI(3,4,5)P_3_ levels, indicating that both PTEN and CnrN are required to maintain the basal PI(3,4,5)P_3_ levels.

AprA increased PI(4,5)P_2_ levels and decreased PI(3,4,5)P_3_ levels in Ax2 cells. AprA alters PI(4,5)P_2_ and PI(3,4,5)P_3_ levels at timepoints when the cells start to respond and migrate away from AprA (Phillips and Gomer, 2012; Rijal et al., 2019), and it is possible that cells move away from the source of AprA by inhibiting PI(3,4,5)P_3_-dependent pseudopod extension at the side of the cells facing toward the source of AprA. Cells lacking PTEN or CnrN do not move away from the source of AprA (Herlihy et al., 2013; Rijal et al., 2019). AprA reduced PI(4,5)P_2_ levels in cells lacking PTEN, but not in cells lacking CnrN, suggesting that loss of PTEN might activate an unknown pathway to cause AprA to decrease PI(4,5)P_2_ levels.

The basal levels of total actin were decreased in *pten^–^* and *cnrN^–^* cells, however, the proportion of F-actin to total actin was increased in both *pten^–^* and *cnrN^–^* cells. This increase in F-actin could be due to the increased levels of PI(3,4,5)P_3_ levels in *pten^–^* and *cnrN^–^* cells, which might be the reasons why *pten^–^* cells have decreased cell migration speed and persistence, and increased basal number of pseudopods and filopods, and PTEN might have compensated the loss of CnrN in *cnrN^–^* cells to regulate cell speed, persistence, and pseudopods (Rijal et al., 2019).

The proportion of polymerized myosin II to total myosin II was increased in both *pten^–^* and *cnrN^–^* cells, possibly due to the increased F-actin accumulation. When the myosin II tail is phosphorylated, the phosphorylated amino acids prevent myosin from forming contractile myosin filaments between the F-actin filaments that allow for the contractile activity at the ‘rear’ of the cell during migration (Levi et al., 2002; Liang et al., 1999; Pasternak et al., 1989). We found that *cnrN^–^* cells had decreased basal phosphorylated myosin levels compared to Ax2 and *pten^–^* cells, suggesting that myosin II is likely to be highly bundled in *cnrN^–^* cells and that CnrN negatively regulates myosin dephosphorylation in *D. discoideum*.

Reduced levels of phosphorylated myosin II and filopod numbers in *cnrN*^–^ cells suggest that PTEN but not CnrN is indispensable for maintaining cell speed and persistence, and both CnrN and PTEN are necessary to maintain basal F-actin and polymerized myosin II levels, and CnrN, but not PTEN, is necessary for maintaining basal phosphorylated myosin II levels. However, loss of CnrN or PTEN abolished the AprA-mediated decrease in phosphorylated myosin, indicating that both CnrN and PTEN are required for AprA to reduce levels of phosphorylated myosin (Rijal et al., 2019).

In a gradient of AprA, *Dictyostelium* cells inhibit Ras activation and pseudopod formation at the side of the cell facing the source of AprA (Kirolos and Gomer, 2022; Rijal et al., 2019). Prolonged exposure (60 minutes) of cells to a uniform concentration of AprA causes cells to become rounder (Kirolos and Gomer, 2022). Loss of PTEN, but not CnrN, caused cells to become rounder in the presence of AprA, suggesting that AprA induces rounding of cells not by inhibiting Ras activation, but by activating a PTEN-dependent unknown pathway. It is possible that AprA decreases the number of pseudopods in *pten*^–^ and *cnrN*^–^ cells after 0 and/or 30 minutes exposure of cells to AprA by Ras independent mechanisms, and the subtle or no effect of AprA on cellular projections such as filopods, pseudopods, and macropinosomes indicates that AprA inhibition of Ras activation causes biased movement of cells not by altering number, size, and lifespan of these projections, but the location of these projections (Kirolos and Gomer, 2022; Rijal et al., 2019) during cell movement.

Our results indicate that CnrN and PTEN are both needed for AprA-induced chemorepulsion, AprA inhibition of proliferation, and AprA inhibition of Ras activation. How two enzymes with similar properties could both be needed for these effects is unclear. One possibility is that neither CnrN nor PTEN is present at sufficiently high levels and thus has enough activity, and for the effects, cells need both activities. Alternatively, although they both localize to the side of the cell closest to the source of AprA in an AprA gradient, CnrN and PTEN might function in different membrane environments (such as small lipid rafts) that cannot be distinguished by optical microscopy. For instance, *D. discoideum* possesses five class I PI3 kinases (Eichinger et al., 2005). PI3 kinase 1/2 produces PI(3,4,5)P_3_ in the membrane domains involved in formation of macropinosome ruffles, whereas PI3 kinase 4 produces PI(3,4,5)P_3_ in the vesicle membrane at the later stage of formation of macropinosomes (Hoeller et al., 2013), and two different Ras GTPases regulate local PI(3,4,5)P_3_ (Hoeller et al., 2013). In support of the idea that CnrN cleaves PI(3,4,5)P_3_ that is present in one type of membrane domain, and PTEN cleaves PI(3,4,5)P_3_ that is present in a different type of membrane domain, we observed that CnrN and PTEN have different effects on basal levels of PI(4,5)P_2_, the ability of AprA to increase the roundness of cells, the number of filopods and pseudopods, and the sizes of filopods. In conclusion, the distinct effects of CnrN and PTEN suggest that these enzymes play different roles in *D. discoideum* signaling pathways, possibly by dephosphorylating PI(3,4,5)P_3_ in different membrane domains to regulate cell responses to AprA.

## Materials and Methods

### Cell strains and culture

*D. discoideum* strains were obtained from the *Dictyostelium* Stock Center (Fey et al., 2019), and were wild-type Ax2, *cnrN^–^* (DBS0302655; (Tang and Gomer, 2008)), *pten^–^* (DBS0236830; (Iijima and Devreotes, 2002)), *pten^–^*/*pten-gfp* (DBS0236831; (Iijima and Devreotes, 2002)), and *cnrN^–^*/*cnrN-gfp* (DBS0302656; (Tang and Gomer, 2008)). Cells were grown at 21°C in shaking culture in HL5 medium (Formedium, Norfolk, England) and on SM/5 agar plates (2 g/L glucose (VWR, Solon, OH), 2 g/L bacto peptone (Becton, Dickinson and Company, Sparks, MD), 0.2 g/L yeast extract (Hardy Diagnostics, Santa Maria, CA), 0.2 g/L MgSO_4_·7H_2_O (Fisher Scientific, Fair Lawn, NJ), 1.9 g/L KH_2_PO_4_ (VWR), 1 g/L K_2_HPO_4_ (VWR), 15 g/L agar (Hardy Diagnostics)) (Sussman, 1966) with a lawn of *Escherichia coli* B/R20 (*Dictyostelium* Stock Center). 100 µg/ml ampicillin and 100 µg/ml dihydrostreptomycin were used to kill *E. coli* in *Dictyostelium* shaking cultures transferred from SM/5 agar plates (Brock and Gomer, 1999). Cells expressing a selectable marker were grown under selection with the appropriate antibiotics (5 µg/ml blasticidin and 5-10 µg/ml G418). GFP expressing cells were grown under constant selection, and the expression of GFP was confirmed by immunofluorescence microscopy.

### Recombinant AprA and chemorepulsion assays

Recombinant AprA was expressed in *E. coli*, purified, stored in 20 mM NaPO_4_ pH 6.2, and checked for purity as described previously (Brock and Gomer, 2005). Chemorepulsion assays were performed using an Insall chamber as previously described (Rijal et al., 2019).

### Fluorescence imaging of fixed cells in an AprA gradient

Imaging of GFP-expressing cells in an AprA gradient was performed as previously described (Rijal et al., 2019). Briefly, *pten^–^/pten-gfp* cells were maintained in log phase (2 to 4 cells/ ml) in HL5 media prior to experiment. 2.4 x 10^4^ cells per well in a volume of 300 µl were allowed to settle in 8-well slides (# 354118, Corning, Big Flat, NY) for 1 hour in a humid chamber. A volume of 1.8 µl of a 50 µg/ml stock of Recombinant AprA in 20 mM NaPO_4_ pH 6.2 was carefully added to the corner of each well and then left to sit undisturbed in the humid chamber for 20 minutes to let the gradient establish. Media removed from the well and 300 µl of 4% paraformaldehyde (# 19210, Electron Microscopy Sciences, Hatfield, PA) in PBS was added to the well and cells were fixed for 10 minutes. The fixative was gently removed, and cells were washed twice with 300 µl of PBS for 5 minutes each and then permeabilized for 5 minutes with PBS/ 0.1% Triton X-100 (# J66624, Alfa Aesar, Ward Hill, MA). Cells were then washed three times for 5 minutes each with PBS and then stained with a 1:3,000 dilution of Phalloidin-555 Alexafluor (# ab176756, Abcam, Cambridge, United Kingdom) in 300 µl PBS for 30 minutes. Cells were washed three times for 5 minutes each with PBS, and then coverslips were mounted with Vectashield hardset mounting medium with 4’,6-diamidino-2-phenylinodole (DAPI) (Vector Laboratories, Burlingame, CA) following the manufacturer’s directions. Images were captured with a 20x objective using a Ti2-Eclipse (Nikon, Kyoto, Japan) inverted fluorescence microscope. Deconvolution of images was done by using the Richardson–Lucy algorithm in NIS-Elements AR software (Laasmaa et al., 2011). Random cells from more than four fields of view in individual experiment were scored by blinded observers as having cytosolic GFP fluorescence, or as having localized GFP fluorescence in the 90° sector of the cell periphery either closest to the AprA source, furthest from the AprA source, or the two the 90° sectors perpendicular to the AprA source. For all microscopy, images of a calibration slide (Swift, Carlsbad, CA) were used to generate size bars and calibrate measurements.

### Proliferation assays

Proliferation of *Dictyostelium* cells in the presence or absence of 300 ng/ml AprA was measured as previously described (Choe et al., 2009), except that the cell density was measured at 24 hours.

### PI(4,5)P_2_ and PI(3,4,5)P_3_ extraction and ELISAs

For the phosphatidylinositol extractions, 1.0 x 10^7^ cells were stimulated with 300 ng/ml AprA for the indicated time. The reaction was stopped with an equal volume of ice cold 1 M Trichloroacetic Acid (TCA) and incubated on ice for 5 minutes. Phosphatidylinositol extractions and ELISAs were performed following the manufacturer’s directions for the phosphatidylinositol 4,5-bisphosphate (PI(4,5)P_2_) Mass ELISA kit (# K-4500) and the phosphatidylinositol 3,4,5-trisphosphate (PI(3,4,5)P_3_) Mass ELISA kit (#K-2500s, Echelon Biosciences Inc, Salt Lake City, UT).

### Reverse transcription (RT-PCR) analysis

The validation of strains was performed as described (Rijal et al., 2019). Briefly, total RNA was extracted from Ax2, *cnrN^–^*, *cnrN^–^/ cnrN-gfp*, *pten^–^,* and *pten^–^/ pten-gfp* strains using a Quick-RNA miniprep kit (# R1054, Zymo Research, Irvine, CA), and cDNA was synthesized using a Maxima H minus first-strand cDNA synthesis kit (# K1652, Thermo Fisher Scientific, Waltham, MA). A PCR reaction was performed to confirm the presence or absence of cDNA from the strains using gene specific primers. *gpdA* primer pair served as a positive control. Oligonucleotides for validating strains by PCR were: *gpdA* forward: 5′-ACCGTTCACGCCATCACTGCC-3′ and reverse: 5′-GACGGACGGTTAAATCGACGACTG-3′; *cnrN* forward: 5′-ACAGGCTTAGAAGCAAGTTGGAGA-3′ and reverse: 5′-ACGTTGTTGTGAAGGTTGAGTTACA-3′; *pten* forward: 5′-AGTTGCAGTCTCTAAACAAAAGAG-3′ and reverse: 5′-GGTGCGTCTGATGCTACAAC-3′. Molecular mass standards for gels were 100 bp and 1 kb DNA ladders (GoldBio, St. Louis, MO).

### Cytoskeletal protein extraction and western blotting

Whole cell actin and myosin II, filamentous actin (F-actin), polymerized myosin II, and phosphorylated myosin II levels were determined exactly as described in (Rijal et al., 2019).

### Cell roundness measurement

For cell roundness measurements, Ax2 cells at 10^5^, 1.5 × 10^5^, 5 × 10^5^, 10^6^, and 5 × 10^6^ cells/ml were cultured in a 96-well, black/clear, tissue-culture-treated, glass-bottom plate (# 353219, Corning) in 300 μl HL5 media. After allowing cells to settle for 30 minutes, images of cells were taken using a 40x objective on a Ti2-Eclipse (Nikon) inverted fluorescence microscope, imaging at least 40 cells per assay. AprA (or an equivalent volume of 20 mM NaPO_4_ pH 6.2) was added to a final concentration of 300 ng/ml. After 30 minutes, cells were imaged as above. The short (As) and long (Al) axes of cells were measured using Fiji (ImageJ) (Schindelin et al., 2012). Roundness was quantified by calculating the ratio of As/Al. To determine if AprA requires CnrN and PTEN to induce roundness, Ax2, *cnrN^–^*, and *pten^–^* cells at 1.5 × 10^5^ cells/ml were assayed as above.

### Ras activation assay

Ras activity in Ax2, *cnrN^–^*, and *pten^–^* cells was assessed using a pull-down assay kit (Cat. #BK008-S; Cytoskeleton, Denver, CO) following the manufacturer’s instructions as previously described (Kirolos and Gomer, 2022).

### Filopods, pseudopods, and macropinosomes quantification

A hole was punched in the bottom of a 100 mm type 25384-302 petri plate (VWR) with a gas flame-heated 13 mm glass test tube. After sanding the resulting burr and washing the plate with distilled water, a 25 x 25 mm glass coverslip was attached to the bottom of the plate covering the hole with heated paraffin wax (Gulf Lite, Memphis, TN). *Dictyostelium* cells from log-phase cultures were washed twice in HL5 medium by centrifugation at 500 x g for 3 minutes and resuspension in 1 ml, and were diluted to 0.15 x 10^6^ cells/ml in 1 ml. A volume of 300 µl of cells was placed on the coverslip in the petri dish and allowed to adhere for 30 minutes. AprA (or an equivalent volume of 20 mM NaPO_4_ pH 6.2) was added to the cells to a final concentration of 300 ng/ml. Images of cells were then captured starting at 0, 30, and 60 minutes in the presence or absence of AprA for 5 minutes, with 2-second intervals, using a 100x oil immersion Hoffman modulation lens ((Modulation Optics, Greenvale, NY) on a Diaphot inverted microscope (Nikon), with the Hoffman condenser in the liquid over the cells to obtain the illumination required for the Hoffman imaging. Images were analyzed using ImageJ (Schneider et al., 2012) to assess the size, lifespan, and count of filopods, pseudopods, and macropinosomes. The ImageJ manual tracking tool was used to measure both the count and lifespan of the structures, the ImageJ freehand selections tool was used to measure the area of pseudopods and macropinosomes, and the straight measure tool was used to measure the size of filopods.

### Statistical analysis

Statistical analyses were performed using Prism 10 (GraphPad Software, Boston, MA) or Microsoft Excel. A p < 0.05 was considered significant.

## Acknowledgements

We thank Dr. Sara Milligan for educational conversations and assisting in experiment design.

## Author Contributions

K.M.C. and R.R. designed and performed experiments, analyzed the data, and wrote the paper; S.B., R.M., K.B., D.C., J.S., and J.S. performed experiments and analyzed the data; and R.H.G. coordinated the study, wrote the paper, and acquired funding. This work was supported by the National Institutes of Health Grant GM118355.

